# Remote spatially variant debiased profiling of cell and tissue mechanical properties

**DOI:** 10.1101/2021.05.12.443111

**Authors:** Jonathan H Mason, Lu Luo, Yvonne Reinwald, Matteo Taffetani, Amelia Hallas-Potts, C Simon Herrington, Vlastimil Srsen, Chih-Jen Lin, Inês A Barroso, Zhihua Zhang, Zhibing Zhang, Anita K. Ghag, Ying Yang, Sarah Waters, Alicia El Haj, Pierre O Bagnaninchi

**Author notes:** Electronic address; Senior and corresponding author. Co-first author.

## Abstract

The role of the mechanical environment in defining tissue function, development and growth has been shown to be fundamental. Assessment of the changes in stiffness of tissue matrices at multiple scales has relied mostly on invasive and often specialist equipment such as AFM or mechanical testing devices poorly suited to the cell culture workflow.In this paper, we have developed a novel unbiased passive optical coherence elastography method, exploiting ambient vibrations in the sample that enables real-time noninvasive quantitative profiling of cells and tissues. We demonstrate a robust method that decouples optical scattering and mechanical properties by actively compensating for scattering associated noise bias and reducing variance. The efficiency for the method to retrieve ground truth is validated in silico and in vitro, and exemplified for key applications such as time course mechanical profiling of bone and cartilage spheroids, tissue engineering cancer models, tissue repair models and single cell. Our method is readily implementable with any commercial optical coherence tomography system without any hardware modifications, and thus offers a breakthrough in tissue mechanical assessment for novel on line assessment of spatial mechanical properties for organoids, soft tissues and tissue engineering.

## Introduction

The mechanical environment in tissue homeostasis has been shown to be fundamental to multiple organ function, development and pathology [1][2, 3]. Matrix stiffness can be an informative indicator in many biological and medical applications. In tissue engineering, the bulk and spatial mechanical properties of engineered grafts is crucial to their clinical success after implantation [4, 5, 6, 7]. For example, nutrient limitation can create a softer central region in engineered cartilage [8, 9]. In cancer research, stiffness differentiates malignant tissue from healthy tissue [10], and monitoring the change in stiffness of 3D cancer cell model in response to anti-cancer drug treatment may potentially indicate the effectiveness of the drug [11]. In the eye, the stiffness of the cornea is indicative of its optical performance under intraocular pressure [12]. Traditional approaches for testing the mechanical properties of engineered tissues usually requires direct contact with the tissue and are non-sterile, involving termination of the cell culture[13, 14]. Furthermore, it provides only bulk values rather than localised insight into the spatial mechanical heterogeneity of the engineered tissue. Manufacturing or long term cultures need easy continuous monitoring without damage to the 3D cultures and optical systems provide a potential solution. Therefore, a system is needed for sterile online monitoring the bulk and spatial mechanical properties of in vitro 3D tissues such as cell seeded matrices, organoids or ex vivo explants

The quantification and spatial mapping of stiffness, a process known as elastography, can be generally performed by stimulating a specimen, measuring its deformation, and inferring its mechanical properties through fitting to a parameterised model. Optical coherence tomography (OCT) [15] is particularly well suited for elastography deformation tracking in small samples, due to its high-resolution non-invasive 3D imaging capacity [16], and its ability to precisely encode displacement through its phase [17].

Early optical coherence elastography (OCE) methods used surface compression with speckle tracking [18, 19] and later phase lag measurement [20], but the concept has been realised with many other contact and non-contact forms of stimulation [21]. One successful approach is to launch controlled shear waves in the material from point dynamic loading via an air-puff [22], and measure the spatially resolved wave velocity using OCT, which is closely tied to the material’s stiffness [23]. Naturally occurring broadband diffuse shear waves can also be exploited to measure shear wavelength [24, 25], a concept used by Nguyen et al. [26] with OCT where it is referred to as ‘passive elastography’. A closely related approach by Zvietcovich et al. [27] measures shear wavelength from reverberant waves from an array of contact point sources vibrating at a single frequency, where it was successfully applied ex-vitro to quantifying cornea stiffness.

Passive elastography is a compelling technique for mechanical contrast, since it can be performed with OCT systems with no additional hardware modifications. Without need for contact with the materials, analysis can be done in a sterile manner for tissue engineering applications. Although the acquisition is both asynchronous and slow relative to the vibrating source, one can still estimate the shear wavelength from a series of displacement fields using the crosscorrelation Bessel fit algorithm of [27] or displacement to strain energy ratio of [26]. However, whilst the former has limited spatial resolution throughout the fitting plane, the latter is inherently biased by varying levels of noise. Therefore, we propose a novel analysis denoted MechAscan for forming a spatially variant debiased estimation of local wavelength, which is suitable for materials of different size and optical scattering properties.

## Results

### MechAscan: robust passive elastography with noise debiasing

First we simulated a shear wave travelling in a biphasic material (Figure 1a) with a wavelength of *λ*1 = 2*mm* in the blue region and *λ*2 = 4*mm* in the yellow region, and tested the performance of our algorithm to retrieve the ground truth with a slow and asynchronous acquisition when compared to the vibrating source. Time-lapse phase measurements (N=195) corresponding to the displacement field induced by the shear wave travelling in the material (x-y plane) are simulated in Figure 1b with an additive white gaussian noise (variance *σ* = 0.05), and used as the inputs to the OCE algorithms estimating the wavelength along the × axis (Figure 1c). The passive OCE algorithm as presented in [26] is biased by noise in the measurements, which we show theoretically in Section 1.3. This effect is demonstrated in Figure 1c, and further investigated with differing noise level variance in Figure 1d. As the noise in displacement estimates from OCT is related to its signal–to–noise ratio [28, 29], this passive elastography contrast is directly tied to the scattering properties of the material. Therefore, materials with identical stiffness but different optical scattering will be incorrectly differentiated with this algorithm. To alleviate this problem, we present a novel approach, detailed in Section 1.3, that actively compensates for this noise bias and reduces the variance through spatial filtering. The effectiveness of this are demonstrated in the wavelength maps in Figure 1a and line profiles in Figure 1c. The debiased estimate corrects for the average error, but introduces a large amount of variance, attributed to the bias–variance trade-off. Finally, our proposed MechAscan algorithm produces a debiased low-variance estimate very close to the ground truth. Most importantly, the trends are consistent for varying nise levels in Figure 1d, which allows the decoupling of scattering intensity and mechanical contrast.

**Figure 1:**
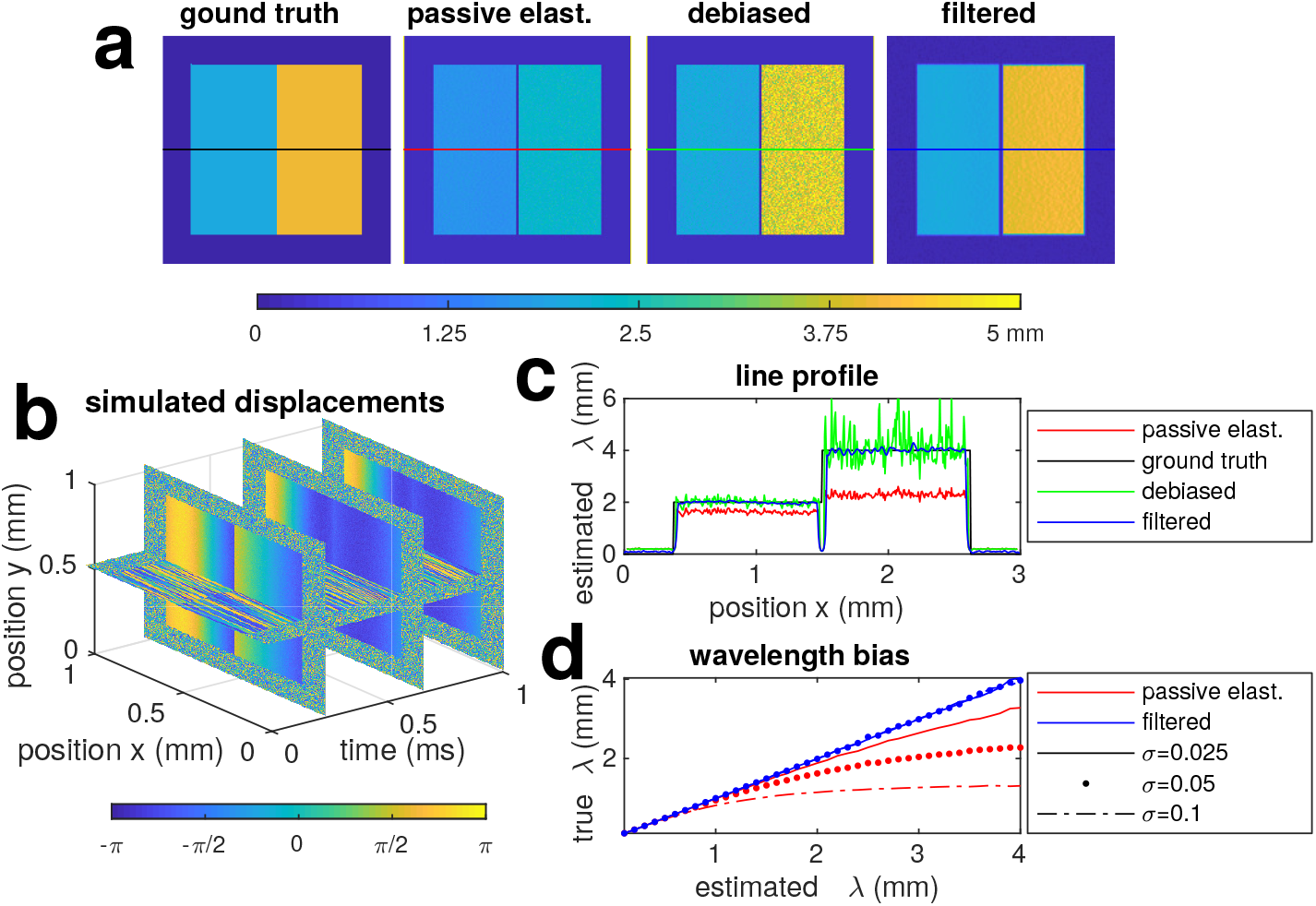
Wavelength estimation from simulation. **a** shows the ground truth with waves travelling laterally in a biphasic material at wavelength *λ*_1_ = 2 mm and *λ*_2_ = 4 mm, the biased estimation from passive elastography as per equation (6) [25], the high variance debiased estimate from equation(8) and the proposed debiased filtered estimate from equation (9), which forms the MechAscan contrast. **b** shows the simulated time-lapse phase measurement induced by the displacement field calculated from ground truth with the addition of a noisy background. **c** shows the line profiles through the coloured regions in **a. d** is the true wavelength against estimated for passive elastography and MechAscan at different noise levels.

Reliably measuring the elastic wavelength then allows one to produce an estimate of stiffness. For linearly elastic isotropic materials for example, *E* ∝ *λ*^2^, where *E* is the Young’s modulus and *λ* is the wavelength estimated through our algorithm. Through calibration, and measurements under the same conditions, the constant of proportionality can be found. More details on the relationship between wavelength and elastic moduli are summarised in Section 1.2.

### Validation with acellular agarose gel

We, then, validated our approach with acellular agarose gels of varying stiffness, which is summarised in Figure 2, with implementation and processing details given in Section 1.1 and experimental protocol in Section 1.4. After OCT imaging, the Young’s modulus of these gels was measured using a standard compression test. The OCT intensity images and calculated mechanical contrast maps of these hydrogels are shown in Figure 2a. All gels had the same amount of optical contrast agent but different concentration of agarose, shown to correlate with its stiffness [30], to decouple optical scattering and mechanical properties. There is a visible increase in stiffness estimate with increasing concentration from the *λ*-maps. Compared to the Young’s modulus of groups of each gel concentration (*n* = 3), as shown in Figure 2c, there is a strong linear correlation (*r* = 0.994) validating the model *E* ∝ *λ*^2^ for this material. We will henceforth refer to the mean wavelength squared, *λ*^2^, as ‘relative stiffness’.

**Figure 2:**
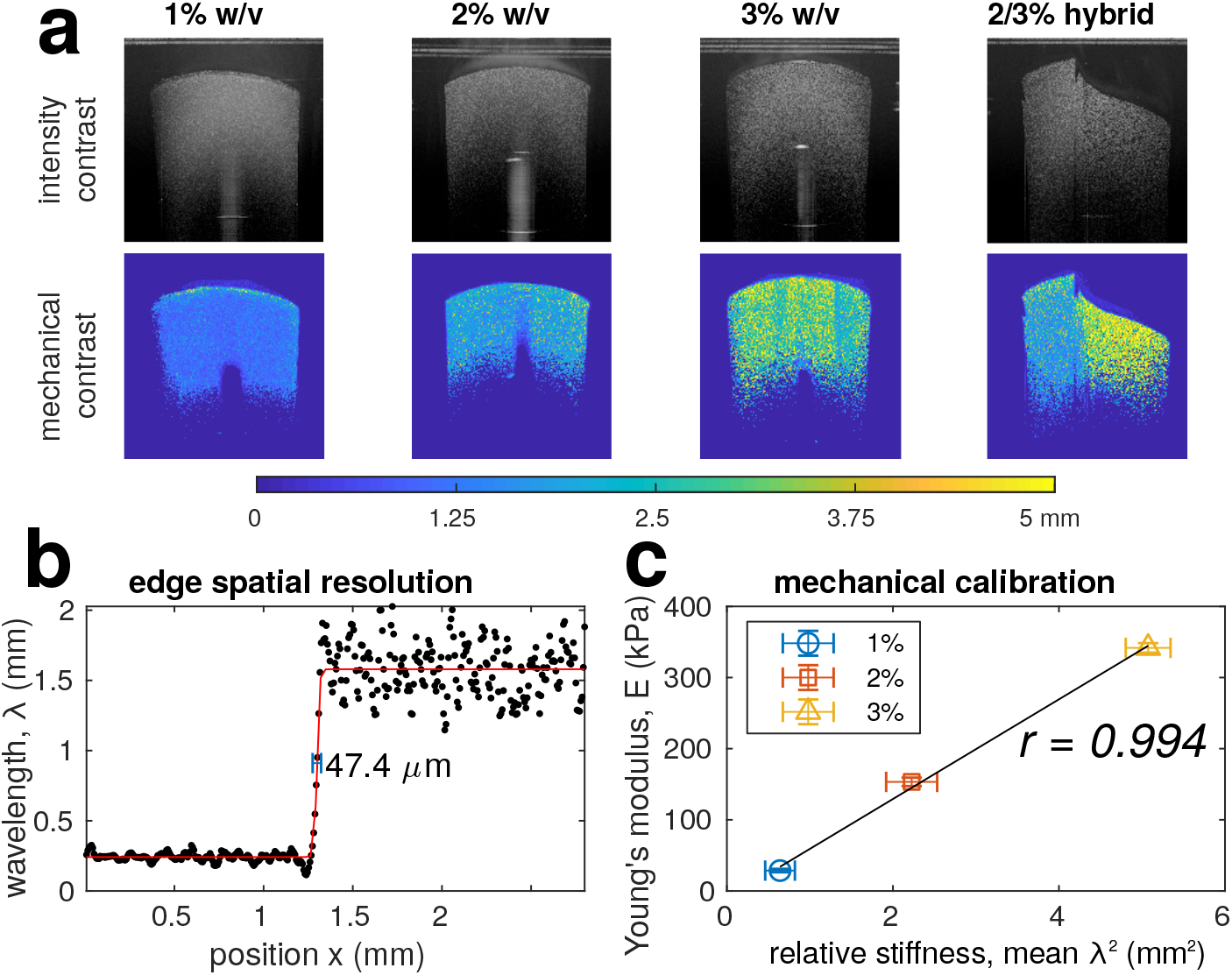
Mechanical contrast and validation with differing concentration agarose gels with 1% w/v milk powder for optical contrast. **a** shows examples of intensity and masked mechanical contrast. **b** is an edge line profile and spatial resolution as calculated as the FWHM of fitted sigmoid derivative as in equation (13); the corresponding axial edge spatial resolution was found to be 38.1*µ*m. **c** mechanical calibration performed as a linear fit between squared wavelength and Young’s modulus from Bose ElectroForce (*n* = 3), assuming the elastic model in equation (3); error bars show the standard deviation. OCT intensity images are displayed as the logarithm of the mean intensity.

The hybrid agarose shown in Figure 2a shows the ability to measure heterogeneous stiffness and local variations, similar to that in [26]. In this case, our algorithm produces a sharp contrast between the two halves of the gel and consistent with depth. Finally, we estimated the spatial resolution of our method and system from the edge transitions, with corresponding profile in Figure 2b, to be 47.4*µ*m laterally and 38.1*µ*m axially, comparable to that reported by [27]; the method to calculate spatial resolution is detailed in Section 1.9.

### On line monitoring mechanical properties of engineered bones and cartilage tissues

Matrix stiffness analysis during orthopaedic tissue engineering is critical to assessment and potentially outcome measures for manufacturing medical implants. As an application of MechAscan analysis in bone tissue engineering, we cultured Mesenchymal Stem Cell (MSC) cell pellets in osteogenic differentiation for 21 days (Section 1.5), while continuously measuring their stiffness with MechAscan in a sterile online manner. Validation was conducted on a subset of sample with end-point mechanical testing. From mechanical testing, shown in Figure 3b, there is a marked increase in the Young’s modulus of engineered bone tissues throughout culture. This increase of mechanical contrast is also clear in the *λ*-maps in Figure 3a in each of the pellets over time. The relative stiffness from MechAscan was then quantified over each of the pellets and shown in Figure 3c, showing a monotonic increase over culture for each sample, in line with the mechanical testing data. Increased staining for collagen in osteogenic pellets was observed with culture time, indicating the growth in matrix content upon culture, further supporting the mechanical testing data and OCT analysis.

**Figure 3:**
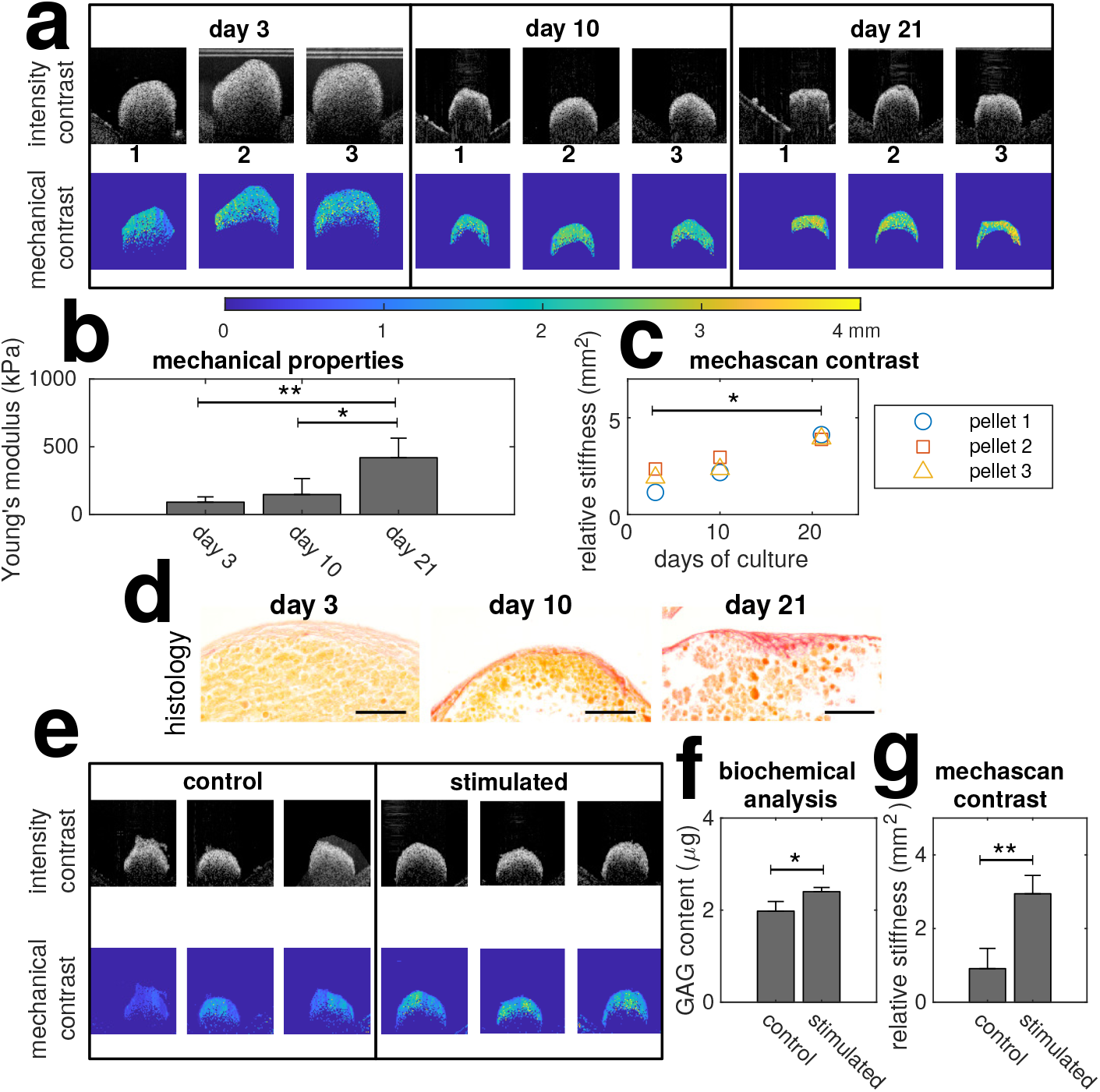
On line monitoring mechanical properties of engineered bones and cartilage tissues. **a** is examples of intensity and masked mechanical contrast of engineered bone tissues (in triplicate) on day 3, 10 and 21. **b** shows the Young’s modulus of the tissue, tested by a customized compression rig (n = 3). [31]. **c** is relative stiffness for for each sample throughout culture (n=3). **d** Histology with picrosirius red staining for collagen content in osteogenic pellets at day 3, 10 and 21. Orange staining indicates cytoplasm and red staining indicates new synthesized collagen fibrils; scale bar: 100 *µ*m. **e** shows intensity and masked mechanical contrast of engineered cartilage tissues stimulated by hydrostatic pressure for 21 days. **f** is glycosaminoglycans (GAG) content of samples (n=3), and **g** shows relative stiffness from MechAscan analysis (n=3).

Then, we applied MechAscan analysis to MSC cell pellets cultured in a chondrogenic medium for 21 days. N=3 samples were subjected to hydrostatic pressure to further enhance the differentiation, while the rest (N=3) were kept as control. As expected, biochemical analysis showed an increase in glycosaminoglycan (GAG) content in the stimulated group as shown in Figure 3f, implying an increase in stiffness, and this is well reflected both in the *λ*-maps in Figure 3e and quantitative analysis summarised in Figure 3g.

### Tissue engineered cancer model

We explored the potential of MechAscan in cancer research for monitoring the spatial mechanical properties of a tissue engineered cancer model. We first cultured fibroblast seeded collagen gels for 11 days. As expected, MechAscan analysis showed a steady increase in the relative stiffness of the fibroblast seeded collagen gel upon culture time Figure 4a,b. Then, ovarian cancer cells were seeded on top of the collagen gel for 7 days, and could easily be distinguished from the collagen gel layer by analysing the mechanical contrast between the two Figure 4c,d.This was supported by histology (Figure 4e,f).

**Figure 4:**
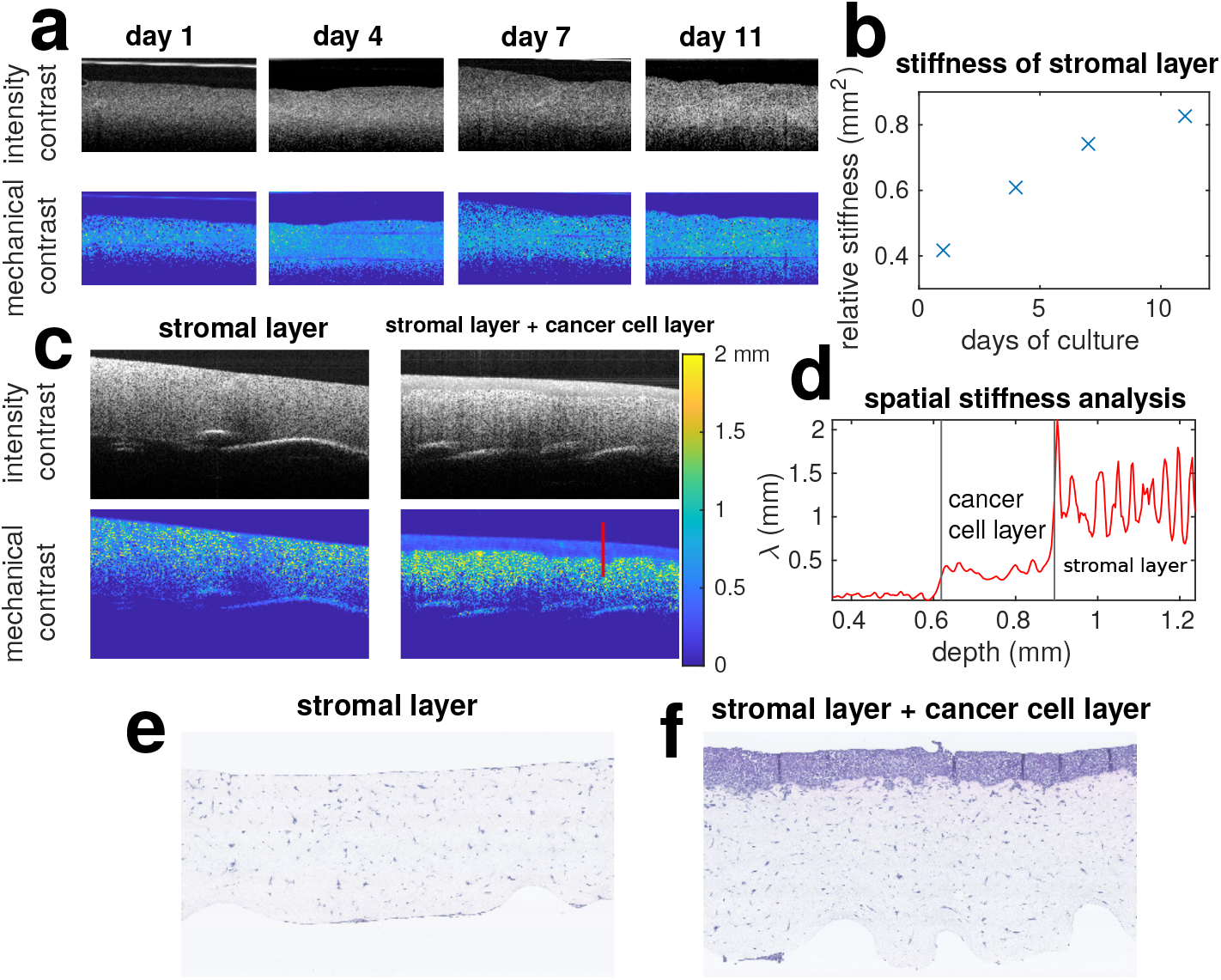
On line mechanical measurement of a tissue engineered cancer model. **a** shows the intensity and mechanical contrast of fibroblast seeded collagen gels over 11 day culture. **b** shows their corresponding stiffness over time. **c** shows the intensity and mechanical contrast of collagen gels with and without cancer cell cultured on top for 7 days. **d** Mechanical properties measured along the red line displayed in **c. e**,**f** Histology images for **c** stained with haematoxylin and eosin.

### Potential application for cornea repair

OCT is a routine tool for ophthalmology and our method could extend these capacities to routine monitoring of surface reconstructions. Thus, we investigated the potential of MechAscan to assess the quality and efficiency of a gel system to repair small cornea lesions. Porcine corneas (N=3) were biopsy punched to generate a local wound. Then, a solution of 15% or 20% methacrylated silk fibroin solution was injected to the injury site, UV crosslinked and imaged with OCT (Fig 5a, Section 1.7). MechAscan analysis showed a significantly higher relative stiffness in the 20% gel group, which is line with the mechanical data obtained by rheology testing (Fig 5 b and c). While OCT intensity images didn’t show any obvious voids at the interface between the gel and corneal in either group, clear voids were identified for both groups in the mechanical contrast (*λ*) map (see white arrows), indicating the existence of uncured gel solution. Burst release pressure for both corneal repair group was found to be 0 kPa (data not shown), further confirming the gel did not fully seal the injury, thus suggesting MechAscan can be a powerful tool for assessing the quality and efficiency of tissue repair.

**Figure 5:**
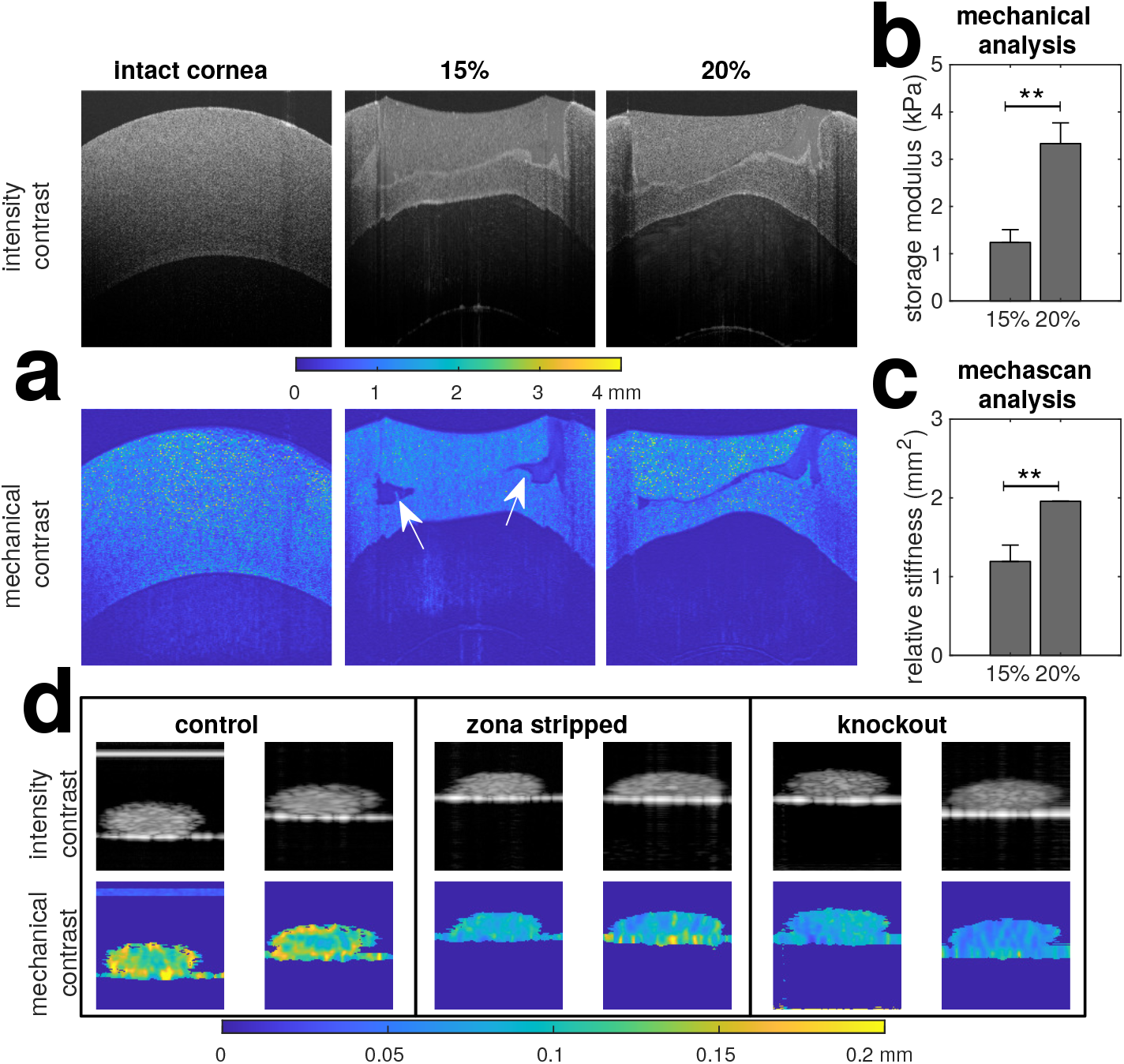
MechAscan applications to tissue samples: **a** shows the intensity and mechanical contrast of porcine corneas intact and after implanting two different concentrations of hydrogel; the white arrows indicate areas of uncured gel, which show high mechanical contrast but not in intensity. **b** shows storage modulus from rheology from samples of hydrogel (*n* = 4). **c** shows relative stiffness from OCT scans. **d** shows a study on the mechanical properties of mouse oocytes: comparing zona intact and zona-free wild type oocytes and intact histone H3.3 knockout oocytes; the zona intact wild type oocytes exhibit greater stiffness heterogenity markedly different from zona-free and less developmentally competent zona-intact H3.3 knockout.

### Potential application for oocyte screening

Routine fertility studies can involve assessment of the viability and developmental competence of pre-fertilized oocytes. Recent studies suggest the stiffness of oocyte is associated with its viability and embryo development potential [32]. The application of the MechAscan analysis to mouse oocytes is summarised in Figure 5d. We compared intact and zona pellucida stripped oocytes from wild-type mice and Cabin1 knockout, which has been shown to cause embryonic developmental arrest [33]. Firstly, a significant difference in mean wavelength was observed against the control group, which could be a means to asses their viability as in [32].Mechanical contrast map showed a stiffer outer ring in wild type zona-intact when compared to zone-free and zona intact cabin1 knockout mice (Fig. 5d). This suggests that MechAscan can be an effective tool for non-contact screening of developmentally competent oocytes in a noninvasive way, in contrast to micropipette aspiration [30].

## Discussion

The MechAscan analysis is a postprocessing algorithm for deriving quantitative mechanical contrast maps from OCT scans, that decouples the scattering and mechanical properties. The approach is based upon the wavelength estimation of ambient elastic waves that are generated by multiple sources in laboratory conditions. Therefore, a necessary condition is the presence of these waves travelling through the sample. From FEM analysis in Section 1.8 we show that external vibrations applied to a well-plate do generate sufficient waves in the sample. Examples of sources include common spectral domain OCT systems as the motorised mirror used to scan through the sample is itself a source of vibration, which can in turn stimulate a vessel such as a well plate if these are mechanically coupled through a stand. The source of these vibrations can also be external environmental from other lab equipment such as refrigeration compressors, incubators or lighting. One must however be conscious that any vibrations must not be excessive, such that they provoke phase wrapping, which introduces bias and artefacts into the estimates. In our experimental setup, we have to take steps to reduce the magnitude of environmental vibrations by placing the plate in a chamber with a rubber base.

Spatial resolution is an important consideration for any imaging method, and the ability to resolve different material stiffness is a main advantage of OCE. Theoretically, our method is limited only by the number of pixels over which the strain power is calculated or the filter kernel. For the agarose study in Figure 2, we calculate strain laterally with a sliding window of 7 pixels and use a Gaussian filter with spatial standard deviation of 5 pixels, leading to a FWHM of 11.8 pixels and expected spatial resolution of 41.3 *µ*m laterally and 20.9 *µ*m axially which is comparable to the 47.4 *µ*m and 38.1 *µ*m we measured empirically from the edge transition. In the presence of noise, there is a trade-off between bias, spatial resolution and variance that can be controlled by the parameters *N*_*g*_ and *N*_*h*_. The higher the noise, the lower a achievable spatial resolution for a given accuracy, so the level of optical scattering is important for high SNR, whilst not attenuating the signal and reducing penetration.

In our study, we have validated the approach using common examples where stiffness is a core assessment during culture or outcome assessment. Our results have shown how we can assess orthopaedic tissues such as cartilage and bone during differentiation in 3D formats, tissue engineered cancer models, fertility assessment of oocytes in vitro, and ophthalmological applications where assessment can be made of biomaterial stiffness with time following implantation.

## Conclusions

MechAscan is a powerful tool for spatially assessing biomaterials and biomatrices in vitro in 3D culture, as it produces a quantitative mechanical contrast image, or *λ*-map, that is decoupled from the standard scattering intensity image from OCT, and this extra information can be used to indicate important changes in the specimen. Additionally, one can extract the ‘relative stiffness’ of a region as the mean squared wavelength, which will be directly proportional to the Young’s modulus of an isotropic linearly elastic material, as validated in agarose samples. Using biological systems with varying matrix properties, we show that this effective and ‘simple to use’ approach offers a breakthrough in tissue mechanical assessment for novel on line assessment of spatial mechanical properties for organoids, soft tissues and tissue engineering.

## 1 Methods

### 1.1 Measurement system and data processing

The system used for the agarose and hMSC pellet study was a Thorlabs Telesto-II spectral domain OCT system with central source wavelength 1310 nm and 243 nm bandwidth. The acquisition consisted of 192 frames, each with 1000 A-scans at a rate of 48 kHz. In the case of the oocyte and cancer seeded collagen gels, the system was a Wasatch Photonics 800 nm system with spectral bandwidth of 92 nm, with measurements consisting of 100 frames of 512 A-scans at a rate of 30 kHz. In all cases, the samples were placed in standard tissue culture well plates on the same bench as the OCT probe to allow transmission of vibrations.

Pre-processing was performed with bespoke software to: resample the spectrometer to k-space, remove the background signal and take the fast Fourier transform to map into complex valued delay space. The displacement fields between subsequent frames were then estimated with

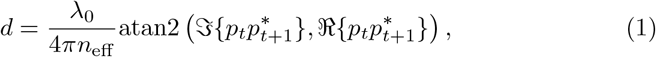

where *λ*_0_ is the central wavelength of the source, *n*_eff_ is the effective refractive index of the sample, ℜ and ℑ take the real and imaginary parts, and *p*_*t*+1_ is the subsequent complex measurement from *p*_*t*_. From these displacement fields, the elastic wavelengths are then estimated through (1.3).

### 1.2 Estimating stiffness from noiseless asynchronous time frames

We wish to estimate the spatially resolved stiffness of a material through its wave speed given as

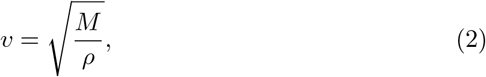

where *ρ* is its mass density and *M* is some elastic modulus factor. This generic form applies to: shear waves, where 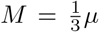, with *µ* the shear modulus; P-waves, where 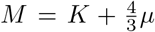, with *K* the bulk modulus; and Rayleigh surface waves, where 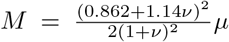 with ν the Poisson’s ratio. As with any travelling wave, the velocity can be expressed as *v* = *λf*, where *λ* is its spatial wavelength, and *f* its temporal frequency. If *f* and *ρ* are either known or known to be fixed a priori, then one can calculate the elastic modulus M directly from measured *λ*.

Assuming shear waves, as in [25, 27], and a linearly elastic isotropic material, the Young’s modulus can be found as

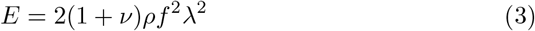

Given periodic elastic waves propagating through a heterogeneous material, one can measure its displacement at a position *x* as

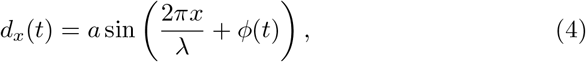

where *a* is the amplitude of the wave, *λ* is the spatial wavelength and *ϕ*(*t*) is a phase function describing the position of the wave at time, *t*. One can also estimate the local strain as a finite difference approximation to a derivative as

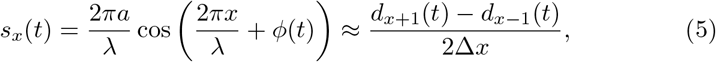

where Δ*x* is the resolution of the imaging system. Given that the energy of a sufficiently sampled sinusoid is *N*_*t*_*a*^2^*/*2 regardless of the order of samples, it follows that *λ* can be found through

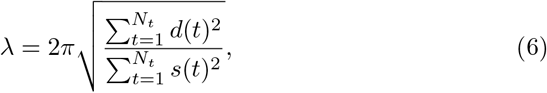

where 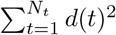 is the displacement energy from a sequence of wave measurements as represented in (4). The estimate in (6) is equivalent to that presented in [25] and used in [26]. The estimate in (6) is exact in a noiseless system as Δ*x* → 0 and the number of temporal samples *N*_*t*_ → ∞. In this case, *ϕ*(*t*) can take the form *ϕ*(*t*) = *kt*, where *k* is a scalar, or a random function spanning [−*π, π*] with uniform probability density. For example, if one has linear sampling with timing jitter such as *ϕ*(*t*) = *kt* + *n* with 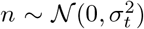, then (6) is still accurate for sufficiently many measurements.

### 1.3 Robust estimation with noise

Although the wavelength estimation in (6) holds in the noiseless case, it is heavily biased by any phase noise in practice. This is because the shear wavelength is in the millimetre range [25, 27], which is far greater than the spatial sampling resolution in OCT, so any additive wide band noise will be amplified by the gradient in (5).

For additive white Gaussian noise (AWGN), *n* ∼𝒩 (0, *σ*^2^), with variance *σ*^2^, the signal energy after filtering with some kernel *h* is

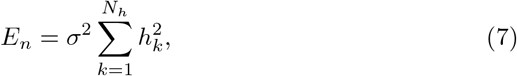

where *N*_*h*_ is the length of *h*. For example, the central difference gradient approximation in (5) is equivalent to spatially filtering the displacement vector with kernel *h* = [− 0.5, 0, 0.5]*/*Δ*x*, which has an energy of *E*_*n*_ = 0.5*σ*^2^*/*Δ*x*^2^. If *σ* can be estimated from the samples, then one can form the debiased estimate of wavelength as

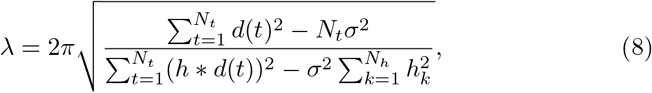

where ∗ denotes spatial convolution.

Whilst (8) may be reasonable in moderately low noise, when the noise energies are similar to the signal energies, it will produce high variance or even undefined estimates. This can be compensated to some degree by increasing *N*_*g*_, but at the expense of spatial resolution in the gradient direction and will not control the numerator going negative. We therefore introduce an additional 2D low-pass filter to the displacement vectors. This debiased filtered estimate as

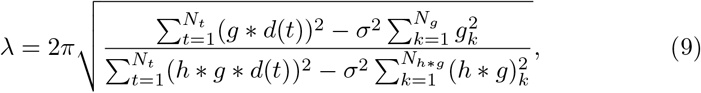

where *g* is the 2D low-pass filter, and *N*_*h*∗*g*_ = *N*_*h*_ + *N*_*g*_ − 1 is the length of the effective strain filter kernel. We use a 2D Gaussian function for *g* since it will not introduce any ripple artefacts, with size *N*_*g*_ = ⌈2*c*_*g*_⌉ + 1, where *c*_*g*_ is its spatial standard deviation of the Gaussian.

Despite the filtering and effective noise power reduction in (9), there still may be cases of undefined estimates due to potential negativity. To avoid this, we adopt a ‘soft subtraction’ approach as used in X-ray computed tomography scatter correction [34]

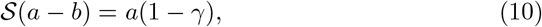

which replaces the operation *a* − *b*, where *γ* is a compensation factor that can be calculated as

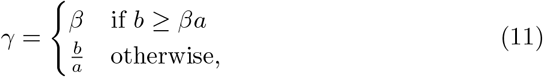

where *β* = [0, 1] is a thresholding term that we set to *β* = 0.9 in this work.

One last consideration for this method is determining the noise variance *σ*^2^. Under the assumptions this is space varying and the wavelength is significantly larger than the spatial resolution of the OCT system, it can be approximated as the time-averaged variance within a sliding 3 *×* 3 window.

### 1.4 Agarose gel study

1%, 2% and 3% agarose hydrogels containing 1% milk were fabricated in this study. Briefly, Type I agarose (Fisher scientific, UK) was added to distilled water and autoclaved at 121 ^°^C to make 2%, 4% and 6% agarose solutions. 2% milk solution was prepared by dissolving nonfat dry milk powder (Bio-rad, UK) in distilled water and this solution was mixed with equal volume of 2%, 4% and 6% agarose solution at 100 ^°^ and cast into a dish and allowed to solidify at room temperature to produce 1%, 2% and 3% agarose gels containing 1% milk. 2/3% hybrid gels were made by scraping part of 2% gel and adding 3% agarose solution into the dish and allowed to solidify. Gels were then cored using a biopsy punch to produce disks of 6mm in diameter and 2 mm in thickness. These gels were then placed into a 96 well tissue culture plate with 100 *µ*l distilled water covering the surface of each gel and imaged with OCT for wavelength measurement as described ealier.

The young’s modulus of these agarose gels were tested using Bose Electro-Force 5500 (TA Instruments, UK). Briefly, excessive water was removed from agarose gels and a preload of 0.01 N was used to ensure direct contact between sample and plate surfaces. An unconfined ramp compression (up to 10% strain of the sample thickness) was performed and the young’s modulus was calculated from the stress-strain curve.

### 1.5 Chondrogenic and osteogenic cell pellet study

Ovine bone marrow derived mesenchymal stem cells were expanded passage two. Cells were then trypsinized and 200,000 cells were added into each well of a v-bottomed 96well plate (Greiner bio-one) and centrifuged for 5 min at 500 g to form cell pellets.

For chondrogenic differentiation, cell pellets were cultured in a differentiation medium consisting of high glucose DMEM supplemented with 2 mM L-glutamine, 100 U/mL penicillin-0.1 mg/ml streptomycin, 100 *µ*g/mL sodium pyruvate, 40 *µ*g/mL L-proline, 50 *µ*g/mL L-ascorbic acid-2-phosphate,

4.7 *µ*g/mL linoleic acid, 1.5 mg/mL bovine serum albumin (BSA), 1 *×* insulin-transferrin-selenium, 100nM dexamethasone (all from Sigma-Aldrich, UK) and 10 ng /ml recombinant Human TGF-*β*3 (Peprotech, UK) for 3 weeks, with medium being exchanged three times a week. During this differentiation period, half of pellets were subjected to hydrostatic pressure (HP, 270 kPa, 1Hz, 1 hour/day) to enhance the differentiation, whereas the rest were not stimulated and kept as control. Both control and HP groups were harvested at end of differentiation period and imaged with OCT for wavelength measurement. Samples were then digested with papain for measurement of their glycosaminoglycan (GAG) content using dimethylmethylene blue dye-binding assay. Samples were also harvested for histology, wax embedded, sectioned and stained with 1% alcian blue 8GX in 0.1 M HCl for GAG distribution.

For osteogenic differentiation, cell pellets were cultured in a differentiation medium consisting of low glucose DMEM supplemented with 10% fetal bovine serum, 2 mM L-glutamine, 100 U/mL penicillin-0.1 mg/ml streptomycin, 100 nM dexamethasone, 50 *µ*M L-ascorbic acid-2-phosphate and 10 mM *β*-glycerco-phosphate (all Sigma-Aldrich, UK) for 3 weeks, with medium being exchanged three times a week. During this differentiation period, samples were continuously monitored using OCT in a sterile manner for wavelength measurement at day 3, 10 and 21, with separate samples in parallel culture being terminated for mechanical testing and histology at same time points. The young’s modulus of osteogenic cell pellets were tested using a customized rig designed for testing microspheres and cell organoids [31]. After excessive water was removed, cell pellets were plated between two platen surfaces and an unconfined ramp compression (up to 25% strain of the sample diameter) was performed and the young’s modulus was calculated from the stress-strain curve. For histological analysis, cell pellets were embedded in wax, sectioned and stained with picrosirius red for collagen accumulation.

### 1.6 Collagen study

Collagen/fibroblast matrices were created using the methodology outlined in Timpson et al [35]. The matrices were incubated at 37 ^°^C, 5% CO2 for 15 days to allow the collagen/fibroblast matrix to contract. The collagen/fibroblast matrix was imaged using OCT on day 1, 4, 7, 11 and 15 of the sample contraction.

Using the protocol outlined in Hallas-Potts et al. [36], one collagen/fibroblast matrix with A2780 cultured on top and one blank collagen/fibroblast matrix with no cells cultured on top were prepared and transferred to a metal grid. Moving the collagen matrix to the grid is referred to day 0. Dishes were incubated at 37 ^°^C with 5% CO2 for 7 days to allow the cells to invade. The collagen/fibroblast matrices were imaged on day 2 and day 7.

Immediately after OCT imaging, the matrix was removed from the grid and added to a falcon tube with 5 mL 4% (w/v) paraformaldehyde (PFA) to fix overnight. The fixed collagen/fibroblast matrix was embedded in wax, sectioned and stained with haematoxylin and eosin (H&E) by the CRUK Edinburgh Centre Pathology & Phenomics Lab. Stained sections were imaged on the Nanozoomer XR model, data captured using NDP scan v 3.1. 40x magnification.

### 1.7 Corneal study

Freshly enucleated porcine eyes were purchased from the local butcher. To induce corneal injury, a 5 mm biopsy punch was used to make a partial cut in the central cornea to a depth of approximately 50%. Then, a solution of methacrylated silk fibroin (15 or 20 wt%) containing 0.5% (w/v) lithium phenyl-2,4,6trimethylbenzoylphosphinate (LAP) was filled to the defect and photocrosslinked for 5 min (365 nm, 3 mW/cm^2^). Fresh corneal, injured corneal and silk fibroin hydrogel repaired corneal was then imaged with OCT for wavelength measurement as described earlier. The storage modulus (G) of freshly prepared Silk-MA hydrogels were measured in oscillatory mode at 32 ^°^C using a plate-plate geometry (Kinexus Pro+, Malvern, UK). After running an amplitude sweep to determine the linear viscoelastic range (LVER), a frequency sweep was conducted at 0.5% shear strain. The experiments were conducted in triplicate.

### 1.8 Finite element method simulation of vibrational elastography

To demonstrate the concept of vibrational elastography, a set of FEM simulations were run, as summarised in Figure 6. This was performed with code aster [37] on a mesh containing a 24 well polystyrene plate and heterogeneous gel with a dynamic load applied axially to one corner of the plate. There were 36000 nodes at the central plane of the gel with thickness 2 mm and diameter 10 mm, whose positions were sampled every 10 *µ*s over a total simulation time of 5 ms. The loading signal used for stimulation was sin(2*πft*)*/t*, where *f* = 2 kHz, with the opposite corner fixed as a boundary condition — other combinations of loads were also tested with similar resulting displacement fields and estimates.

**Figure 6:**
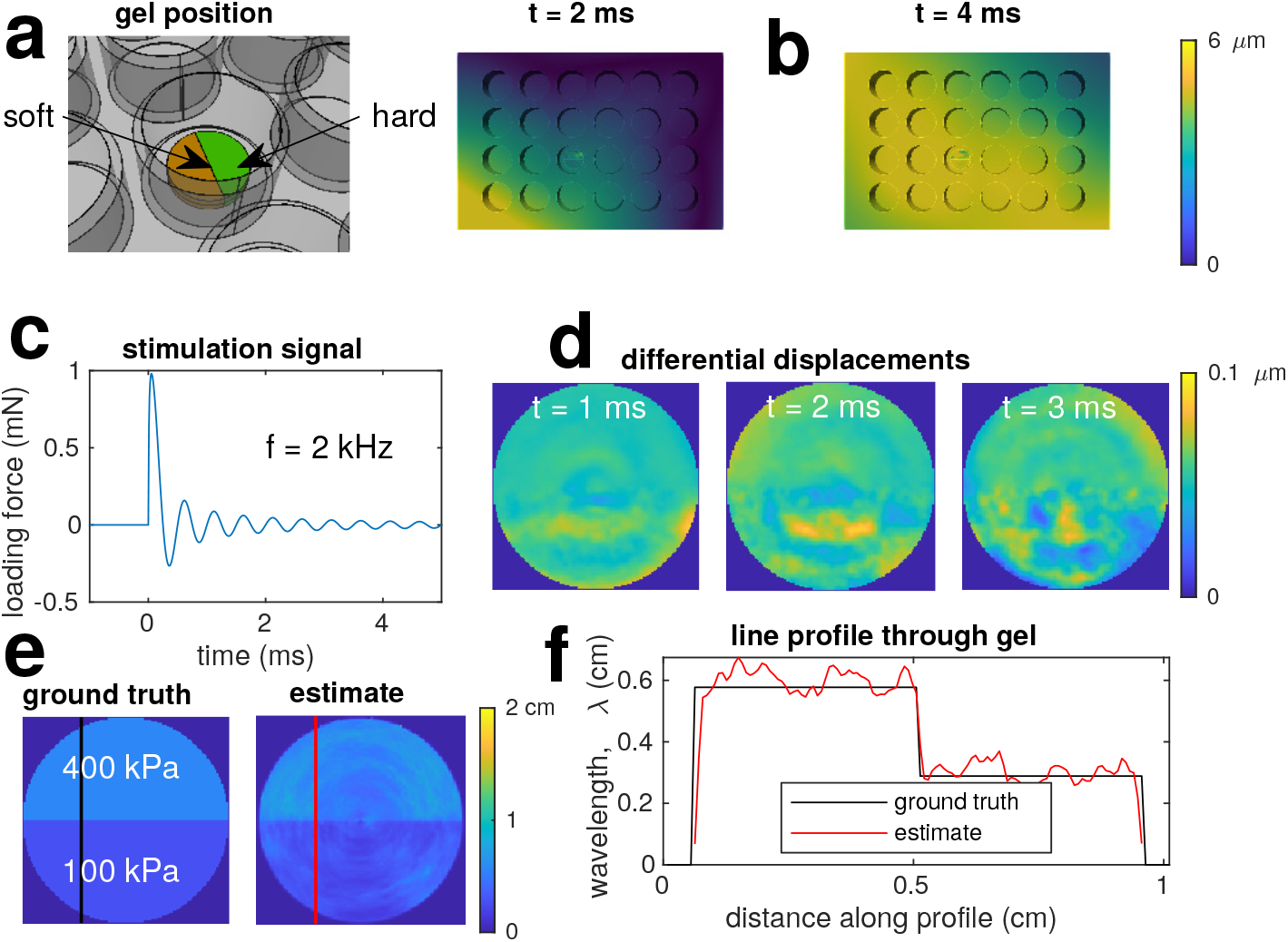
FEM simulation of vibrating multi-well plate with heterogeneous gel. **a** shows the gel seated on the base on a well. **b** shows the displacement magnitudes of the plate at t = 2 ms and t = 4 ms. **c** shows the excitation force applied to a corner of the plate: a 2 kHz sinusoid with exponentially decaying amplitude. **d** shows exampled of the differential displacements in the axial direction at the centre of the gel. **e** shows the estimated wavelength over the gel surface and the ground truth assuming shear waves of 2 kHz. **f** are profile plots are the lines indicated in **e**.

The wavelengths of any resulting elastic waves in this noiseless case were estimated using (6). Firstly, the axial positions at each time instance were resampled onto a polar grid with origin at the centre of the gel so that the strains were estimated along effective cross-sections as used in the experimental section. The estimate from (6) was applied to the 400 displacement (differential position) fields between *t* = [1, 4] ms. Finally, these wavelengths were resampled onto a Cartesian grid to form an image.

### 1.9 Determining spatial resolution

We use a Blonski [38] edge method variant also adopted by [27] for the spatial resolution: first fitting the sigmoid function

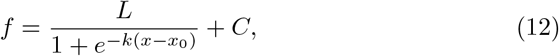

with the Levenberg–Marquardt algorithm for the parameters (*L, k, x*_0_, *C*) then finding the full-width at half-maximum (FWHM) of the sigmoid derivative, which can be expressed as

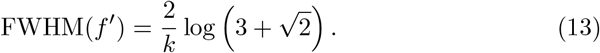

## Notes

### Competing Interest Statement

The authors have declared no competing interest.

